# Cellular uptake of micro and nano plastics Induces Mitochondrial Dysfunction

**DOI:** 10.64898/2026.06.24.734306

**Authors:** Pankhi Vatsa, Vasanthi Rajasekaran, Shubham Dubey, Pulin Che, Yajing Wang, Dan E Berkowitz, Praveen K Dubey

**Author notes:** **Correspondence address: Praveen Kumar Dubey, Ph.D**. Department of Anesthesiology and Perioperative Medicine, School of Medicine, The University of Alabama at Birmingham 901 19^th^ Street S., Birmingham, AL 35205.

## Abstract

Micro and nano plastics (MNPs) have become ubiquitous contaminants in the environment with their occurrence being detected in air, water and food. They can cross biological barriers and slowly build up in different organs, including the placenta, raising concerns about possible impacts on maternal and fetal health. Placenta, a highly metabolically active organ composed primarily of trophoblast cells, requires substantial energy for proper development and function. However, the effects of MNPs exposure on trophoblast biology and mitochondrial health remain poorly understood. This study investigated the in vivo systemic accumulation of MNP in different organs of pregnant mice and their localization within various organelles in vitro. These effects influenced trophoblast energy metabolism and led to reduced migration. Mice received fluorescent polystyrene MNPs via their drinking water. Biodistribution was evaluated in vivo using IVIS whole-body imaging, while ex vivo fluorescence imaging confirmed accumulation of these particles in multiple organs and cells. In parallel, human HTR-8/SVneo trophoblast cells were exposed to MNPs, demonstrating rapid cellular uptake and mitochondrial and nuclear localization via fluorescence microscopy. TEM analysis uncovered mitochondrial structural alterations and the localization of MNPs. Seahorse analysis revealed impaired mitochondrial respiration and oxygen consumption rates, indicating compromised cellular bioenergetics in MNPs-treated cells, which led to inflammation, altered mtDNA copy number, and impaired trophoblast migration. Overall, these findings indicate that pregnant mice exposed to MNPs undergo systemic transfer, with trophoblast uptake marked by mitochondrial dysfunction, inflammation, and reduced migration. Our study identifies mitochondrial dysfunction as a central mechanism underlying MNP-mediated placental toxicity and underscores the potential role of environmental microplastic exposure in adverse pregnancy outcomes.

**Highlights:**

- Cellular uptake of MNPs and their localization across various organelles.
- Uptake of MNPs disrupts mitochondrial function and promotes inflammation.
- Cellular uptake of MNPs impairs cell migration.

## Introduction

Micro and nano plastics (MNPs) are ubiquitous environmental pollutants generated through plastic manufacturing, subsequent use, and environmental degradation [1]. Due to their widespread environmental presence and constant human exposure, MNPs pose a potential public health risk [1–3]. Emerging evidence indicates their presence in various human tissues, including the heart, lungs, kidneys, placenta, breast milk, umbilical cord blood, and maternal circulation. This indicates direct maternal-fetal exposure and an elevated risk of birth complications [4–9]. Placental accumulation of specific polymers (e.g., polyethylene, polystyrene) is associated with elevated preterm birth risk in humans [10,11]. Consistently, animal studies show that maternal exposure to MNPs leads to adverse pregnancy outcomes, including preterm birth, miscarriage, and fetal growth restriction. [5,12–14].

Placenta, a highly metabolically active organ essential for fetal development, consists of trophoblasts that differentiate after egg fertilization [15,16]. These processes enable the efficient transfer of nutrients and oxygen from the mother to the fetus [17]. Disruption of trophoblast function is closely associated with the development and progression of preeclampsia (PE), a hypertensive pregnancy disorder characterized by new-onset hypertension and proteinuria after 20 weeks of gestation[7]. Impaired trophoblast invasion in PE triggers placental ischemia, reactive oxygen species (ROS) overproduction, oxidative stress, endothelial dysfunction, and systemic inflammation, culminating in placental insufficiency and adverse maternal fetal outcomes [17,18]. Mitochondria are central regulators of trophoblast physiology and placental development, governing ATP production, redox homeostasis, and cellular differentiation. Mitochondrial dysfunction has been associated with pregnancy-related complications, particularly preeclampsia, which is characterized by heightened oxidative stress, impaired trophoblast function, and inflammation [19–21]. These mitochondrial abnormalities contribute to placental insufficiency and adverse maternal and fetal outcomes [19,20,22].

In the present study, we are investigating the cellular uptake and biodistribution of MNPs in pregnant mice and human trophoblasts. We hypothesize that the accumulation of MNPs in various organs impairs their function. The internalized MNPs disrupt mitochondrial bioenergetics, leading to increased oxidative stress and inflammatory responses. Ultimately, these effects may compromise trophoblast function and contribute to the pathophysiology of preeclampsia.

## Material and Method

### Vertebrate animals

The University of Alabama at Birmingham’s Institutional Animal Care and Use Committee (IACUC #23159) approved all animal experiment protocols. We used adult mice (10-12 weeks) and housed them in a pathogen-free animal facility.

### IVIS Imaging

For IVIS imaging, green (G25) or red (R25) labeled polystyrene particles (PS) were administered orally (Drinking water) or intraperitoneally (injection) to adult mice. Whole-body live animal imaging was performed 24 hours after intraperitoneal (IP) injection and 72 hours after oral administration. This was done using the IVIS Lumina III imaging system at the University of Alabama at Birmingham’s Small Animal Imaging Facility (SAIF) to evaluate the systemic distribution and accumulation of fluorescent MNPs in vivo.

### Systemic distribution of fluorescence MNPs in pregnant mice

The Adult mice received MNPs (Green, G25, 50 µg/mL) in drinking water, resulting in a concentration of approximately 10 mg/kg/day for two weeks, and then were bred and maintained at the same dose during pregnancy. At E18.5, organs were collected and fixed for imaging.

### Cell culture and treatment

Human trophoblast cells (HTR-8/SVneo; ATCC, Catalog No. CRL-3271) were purchased from the American Type Culture Collection (ATCC, USA) and cultured according to ATCC protocols. They were maintained in RPMI-1640 medium (ATCC, Catalog No. 30-2001) supplemented with 5% fetal bovine serum (FBS; HyClone, Catalog No. SH30028FS) and 1% penicillin-streptomycin (Thermo Fisher Scientific, Catalog No. 10378016) in a humidified incubator at 37°C with 5% CO_₂_ under standard sterile conditions. Depending on the experimental requirements, cells were seeded in 24-well plates, 6-well plates, or 10 cm culture dishes and grown to approximately 70-80% confluence prior to treatment. Trophoblast cells were then exposed to 50 µg/mL MNPs for 24 or 96 h. Following exposure, cells were harvested for RNA and DNA isolation, transmission electron microscopy (TEM), wound-healing assays, and western blot analysis to assess MNP-induced ultrastructural and molecular alterations.

### Cellular uptake of green and red MNPs by trophoblast cells

Intracellular uptake of micro and nano plastics by trophoblast cells was assessed by immunofluorescence staining. HTR-8/SVneo cells were seeded onto sterile glass coverslips precoated with 0.02% gelatin (Catalog No. G9391, Sigma-Aldrich, USA) and fibronectin (1 mg/mL; Catalog No. F1141, Sigma-Aldrich, USA) and allowed to adhere for overnight under standard culture conditions (37°C, 5% CO_₂_). Cells were then exposed to fluorescently labeled red and green polystyrene particles (25 nm; R25 or G25 respectively, Fisher Scientific, USA) for 24 h. After treatment, cells were washed with PBS to eliminate non-internalized particles, fixed with 4% paraformaldehyde for 10 minutes, permeabilized with 0.5% Triton X-100, and blocked with 0.1% BSA to minimize nonspecific binding.

For organelle and structural visualization, mitochondria were stained using primary antibody against TOM20, followed by incubation with the appropriate fluorophore-conjugated secondary antibody. The nuclear envelope was labeled with Lamin B1 antibody. The actin cytoskeleton was visualized using phalloidin staining. Coverslips were mounted with antifade mounting medium with DAPI and imaged using an ECHO Revolve fluorescence microscope. Co-localization analysis was performed to determine the intracellular distribution of fluorescent particles relative to mitochondria, the nucleus, and cytoskeletal structures.

### Cellular Bioenergetics

Cellular bioenergetics were assessed in control and MNP-treated HTR-8/SVneo trophoblasts cells using Seahorse XF Pro M FluxPak (#103775-100) and the Agilent Technologies Seahorse XF Pro Analyzer. Trophoblast cells were plated on Seahorse XF Pro M cell culture 96-well plates and allowed to attach overnight under standard culture conditions. Following attachment, cells were exposed to MNPs at a concentration of 50 µg/mL for 24 h and 96 h, while control cells were maintained under identical conditions served as the control group. After treatment, mitochondrial respiration and metabolic function were analyzed using the Seahorse XF Cell Mito Stress Test protocol as per the manufacturer instructions. Prior to the assay, the culture medium was replaced with Seahorse XF assay medium, supplemented as recommended by the manufacturer, and cells were incubated in a non-CO_₂_incubator to equilibrate. Key mitochondrial parameters, including basal respiration, ATP-linked respiration, proton leak, maximal respiratory capacity, spare respiratory capacity, and non-mitochondrial respiration were evaluated following sequential injections of Oligomycin A (#75351-5MG), FCCP (#75351-5MG), and rotenone (#557368)/antimycin A (#A8674) from Sigma. Oxygen consumption rate (OCR) measurements were recorded using the Seahorse XFp Analyzer to assess mitochondrial function and cellular metabolic activity in both control and MNP treated trophoblast cells. Data acquisition and analysis were performed using ABI WavePro software.

### Transmission Electron Microscopy

Ultrastructural analysis of trophoblast cells was performed using transmission electron microscopy (TEM). Following experimental treatments, trophoblast cells were fixed in 2.5% glutaraldehyde prepared in 0.1M cacodylate (pH 7.4) at 4°C overnight. After primary fixation, samples were washed and post-fixed in 1% osmium tetroxide for 1-2 h at room temperature. The samples were subsequently dehydrated through a graded ethanol series followed by propylene oxide treatment and embedded in epoxy resin. Ultrathin sections (approximately 60-90 nm) were prepared using an ultramicrotome and mounted onto copper grids. Sections were stained with uranyl acetate and lead citrate to enhance contrast and examined using a transmission electron microscope operated at appropriate accelerating voltage settings. Representative images were acquired to evaluate cellular and subcellular ultrastructural alterations, including mitochondrial morphology, nuclear integrity, membrane damage, vacuolization, and the intracellular localization of microplastic particles in trophoblast cells. All samples were imaged on a TEM (JEOL 1400 HC Flash electron microscope with an AMT digital camera). Electron microscopy was performed at our core facility: the UAB High Resolution Imaging Facility.

### Mitochondrial DNA Copy Number Analysis

Genomic DNA was extracted from control and MNP treated trophoblast cells using a commercial DNA extraction kit, Qiagen DNeasy Blood & Tissue Kit (Catalog No. # 69504), according to the manufacturer’s instructions. The quality and quantity of each DNA sample were measured using a NanoDrop spectrophotometer (Thermo Fisher). Mitochondrial DNA (mtDNA) copy number was quantified by quantitative real-time PCR (qPCR) using mitochondrial-specific primers and a nuclear gene (Supplementary table). Relative mtDNA copy number was determined by amplifying a mitochondrial target gene and normalizing its expression to that of a nuclear reference gene. qPCR reactions were were performed using PowerUp™ SYBR™ Green Master Mix (#A25778, Thermofisher). Relative mtDNA copy number was calculated using the comparative C_t_ (2^-ΔΔCt^) method based on the C_t_ differences between mitochondrial and nuclear genes. All samples were analyzed in triplicate, and the results were expressed relative to the control group.

### RNA isolation and qPCR

RNA was isolated with a Qiagen RNeasy Mini Kit, and cDNA synthesis was performed using a Thermo Fisher Scientific reverse transcription kit following the manufacturers’ instructions. Quantitative real-time PCR (qRT-PCR) was performed to measure the expression of GAPDH, TLR9, NLRP3, and IL1β. Amplification was carried out on a QuantStudio 3 Real-Time PCR System (Thermo Fisher Scientific) using PowerUp Universal SYBR Green Master Mix (Life Technologies). Gene expression levels were normalized to the housekeeping gene GAPDH. The PCR cycling conditions were as follows: 50°C for 2 min, 95°C for 3 min, followed by 40 cycles of 95°C for 10 s and 60°C for 30 s. Relative gene expression was calculated using the 2^−ΔΔCt^ method, with vehicle treated samples used as the reference control.

### Wound healing

Cell migration was assessed using the wound healing assay. Briefly, trophoblast cells were seeded in a 6-well plate and treated with MNPs overnight. Further, the cells were replated onto a 24-well plate under above conditions until approximately 90-100% confluency was achieved. A uniform scratch was created across the cell monolayer using sterile 200 µL tip. The detached cells and debris were gently removed by washing with 1 X phosphate-buffered saline (1 X PBS). Images of the scratched area were captured immediately after scratching (0 h) using an inverted phase-contrast microscope. Cells were further incubated at 37°C with 5% CO_₂_, and wound closure was monitored at designated time points (0 and 72h). Images were acquired from the same regions at each time point to assess cell migration into the scratched area. The wound area was quantified using ImageJ software, and the percentage of wound closure was calculated using the following formula: Percentage of Wound closure = Initial wound area at 0 h (A_0_)- wound area at 72h (A_t_)÷Initial wound area at 0 h.

### Statistical analysis

The data were analyzed with a two-tailed unpaired t-test for comparing two groups or using one-way ANOVA after the Brown-Forsythe test and Bartlett’s corrected statistic. All values are shown as mean ± SEM. A P-value of ≤0.05 was considered statistically significant.

## Results

### Biodistribution of radiolabeled microplastics

To assess the biodistribution and accumulation of micro and nano plastics (MNPs) following exposure, in vivo fluorescence imaging was performed using an in vivo imaging system (IVIS) at 24 h (red-labeled MNPs) and 72 h (green-labeled MNPs) post-administration. Vehicle-treated control mice exhibited negligible background fluorescence (Data not shown), whereas MNP-treated mice displayed distinct red and green fluorescence signals distributed throughout the body at the respective time points (Figure 1A, C). These observations indicate systemic accumulation and biodistribution of MNPs following exposure. To further confirm cellular uptake and internalization of green-labeled MNPs, peritoneal cells were harvested 72 h after injection. Cells were seeded onto coated glass slides and incubated overnight to allow adherence. Subsequently, cells were washed, fixed, and stained with wheat germ agglutinin (WGA)-red to label the plasma membrane and counterstained with DAPI to visualize nuclei (Figure 1B). Fluorescence microscopy revealed intracellular green fluorescence within peritoneal cells, confirming successful uptake and internalization of MNPs.

**Figure 1:**
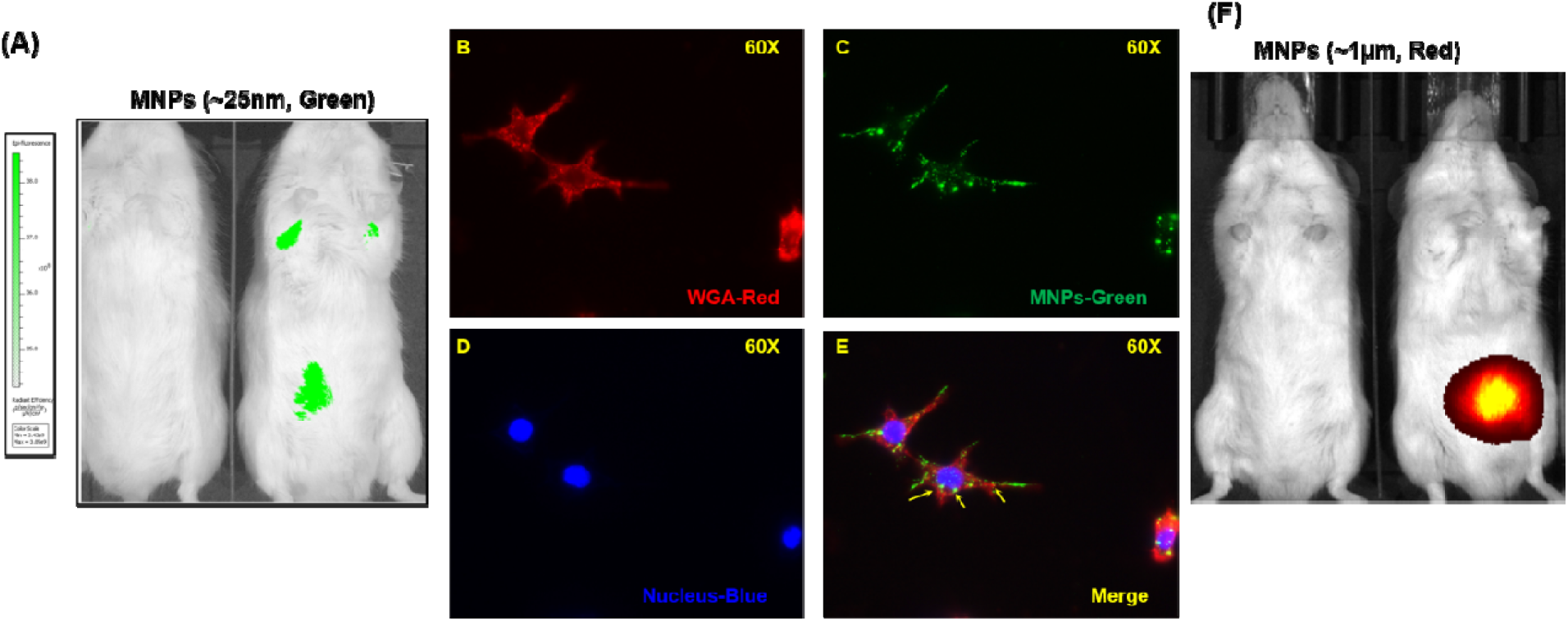
Uptake and biodistribution of polystyrene MNPs in mice. (A) In vivo IVIS fluorescence images of live mice 72 hours after oral administration of 25 nm green fluorescent polystyrene micro and nano plastic particles (MNPs) show their whole-body biodistribution. (B–E) Fluorescence microscopy of peritoneal cells isolated from same mice post 72 h post-MNP exposure, demonstrating cellular uptake of MNPs. (F) Cell membranes stained with wheat germ agglutinin (WGA, red). (C) Internalized MNPs (green). (D) Nuclei counterstained with DAPI (blue). (E) Merged image showing intracellular localization of MNPs (yellow arrows). Images were acquired at 60X magnification. (F) Representative *in vivo* IVIS fluorescence images of live mice 24 h after intraperitoneal injection of red fluorescent polystyrene (1µm). Data are presented as mean ± SEM (*n* = 4).

### Systemic distribution of fluorescent MNPs in pregnant mice

Systemic distribution and placental accumulation of MNPs in mouse tissues and organs were assessed using fluorescence microscopy on maternal organs harvested at embryonic day 18.5 (E18.5). Female mice were orally exposed to green fluorescent 25 nm polystyrene (PS) MNPs for two weeks prior to breeding and continuously throughout gestation. At E18.5, organs were collected, fixed, and imaged. Green fluorescent particle signals (red arrows) were detected in the spleen, kidney, heart, liver, lung and placenta, confirming MNP presence across these tissues (Figure 2). Placental MNP detection indicates uptake and retention, providing direct visual evidence of translocation across the maternal fetal interface and potential fetal exposure. These data demonstrate systemic absorption and multi-organ accumulation of MNPs under environmentally relevant oral exposure conditions.

**Figure 2.**
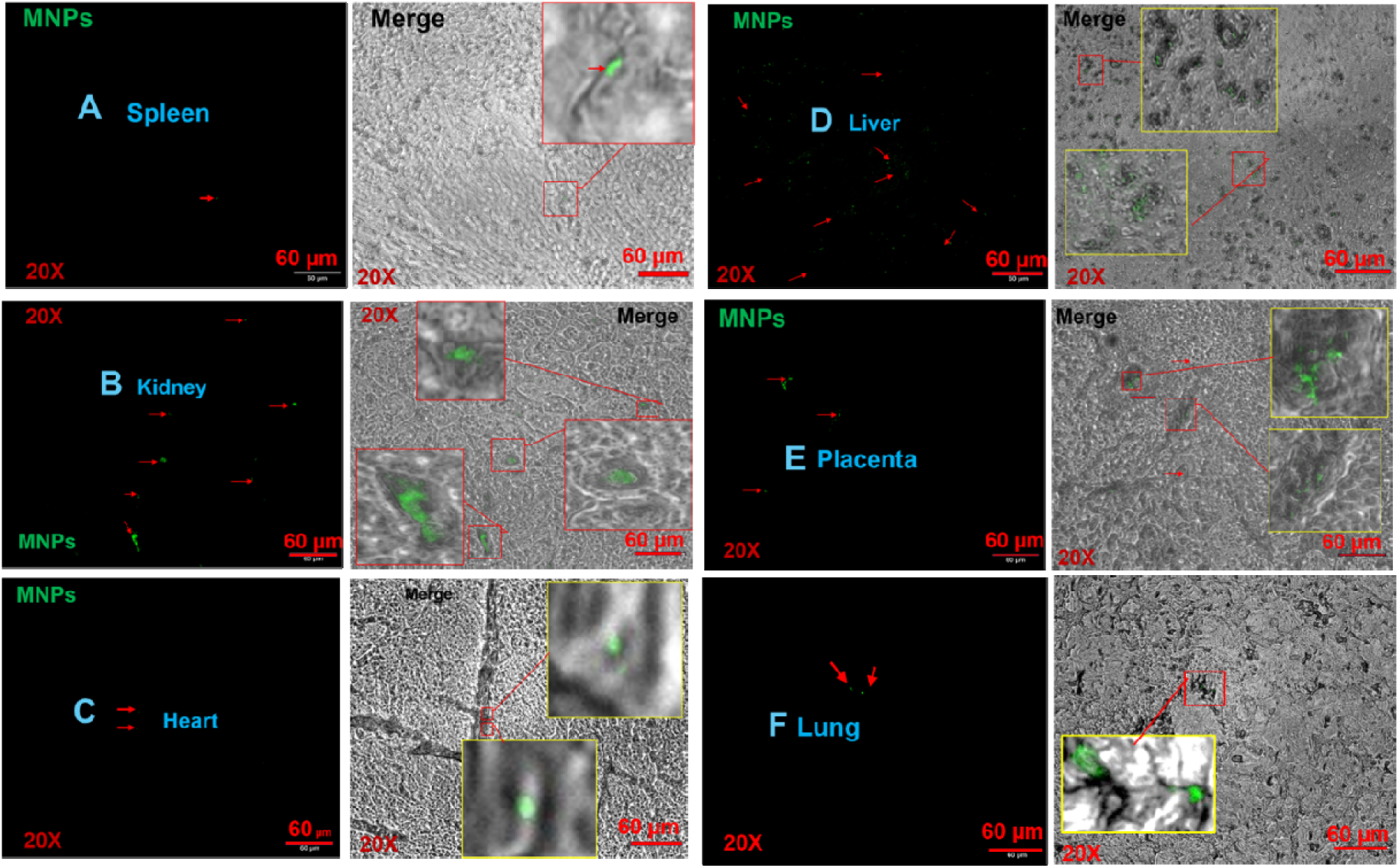
Tissue distribution of green fluorescent polystyrene micro and nano plastics (MNPs) following oral exposure. Fluorescence and brightfield images show green fluorescent Polystyrene micro and nano particles (MNPs) in maternal tissues collected at E18.5 after MNP exposure. (A) spleen, (B) kidney, (C) heart, (D) liver, (E) placenta, and (F) Lung. Green signals indicate MNPs, and arrows highlight representative particles or aggregates within tissue sections. Merged images confirm the localization of MNP fluorescence within tissue structures. Boxed regions are shown at higher magnification to illustrate intracellular or perivascular localization. All images were acquired at 20X. Scale bars = 60 μm.

### Uptake and Subcellular Localization of MNPs in Trophoblast Cells

To investigate the uptake and localization of MNPs in human trophoblast cells, HTR-8/SVneo cells were treated with fluorescently labeled polystyrene (PS) MNPs (green and red). Fluorescence microscopy revealed efficient intracellular uptake of the green PS-MNPs within the cytoplasm and significant accumulation in the nucleus, co-localizing with DAPI-stained nuclei. Similarly, red-labeled MNPs were also found in the nucleus adjacent to the Lamin B1-positive nuclear envelope after 24 hours (Figure 3A, B, C). Furthermore, green MNPs showed spatial overlap with the mitochondrial protein TOM20, indicating an association with mitochondria and suggesting proximity to energy-demanding compartments (Figure 4A). To assess ultrastructural effects, transmission electron microscopy (TEM) was performed. Vehicle-treated cells exhibited normal, well-structured mitochondria, while MNPs treated cells showed rounded mitochondria with disorganized cristae and electron-dense MNP particles (Yellow Arrow) nearby, indicating mitochondrial stress from MNP exposure (Figure 4B). These findings highlight efficient MNPs internalization in trophoblast cells, their close association with mitochondria, and structural alterations, suggesting a potential impact on cellular function and placental health. These findings underscore the need to validate mitochondrial and nuclear intercalation of MNPs and encourages further exploration.

**Figure 3.**
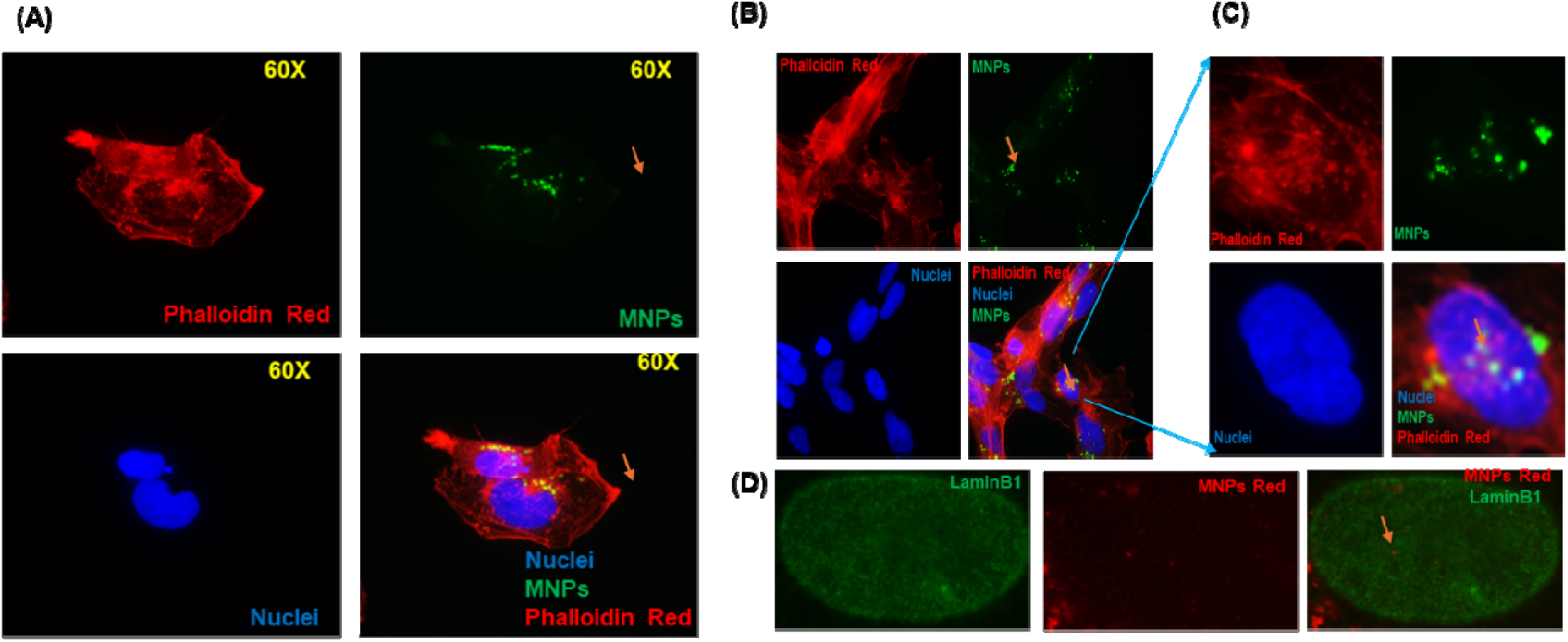
Showing cellular uptake and organellar localization of MNPs by trophoblast cells. (A) Green MNPs (25nm) internalized by trophoblast stained with phalloidin-red staining for the actin cytoskeleton and DAPI (blue) for nuclei. Green fluorescent MNPs are observed in the cytoplasm, indicating their internalization. (B) Green MNPs cluster near and inside the nucleus (Blue). (D) Immunofluorescence with Lamin B1 (green) and red fluorescent MNPs (25nm) showing MNPs within the nuclear envelope.

**Figure 4.**
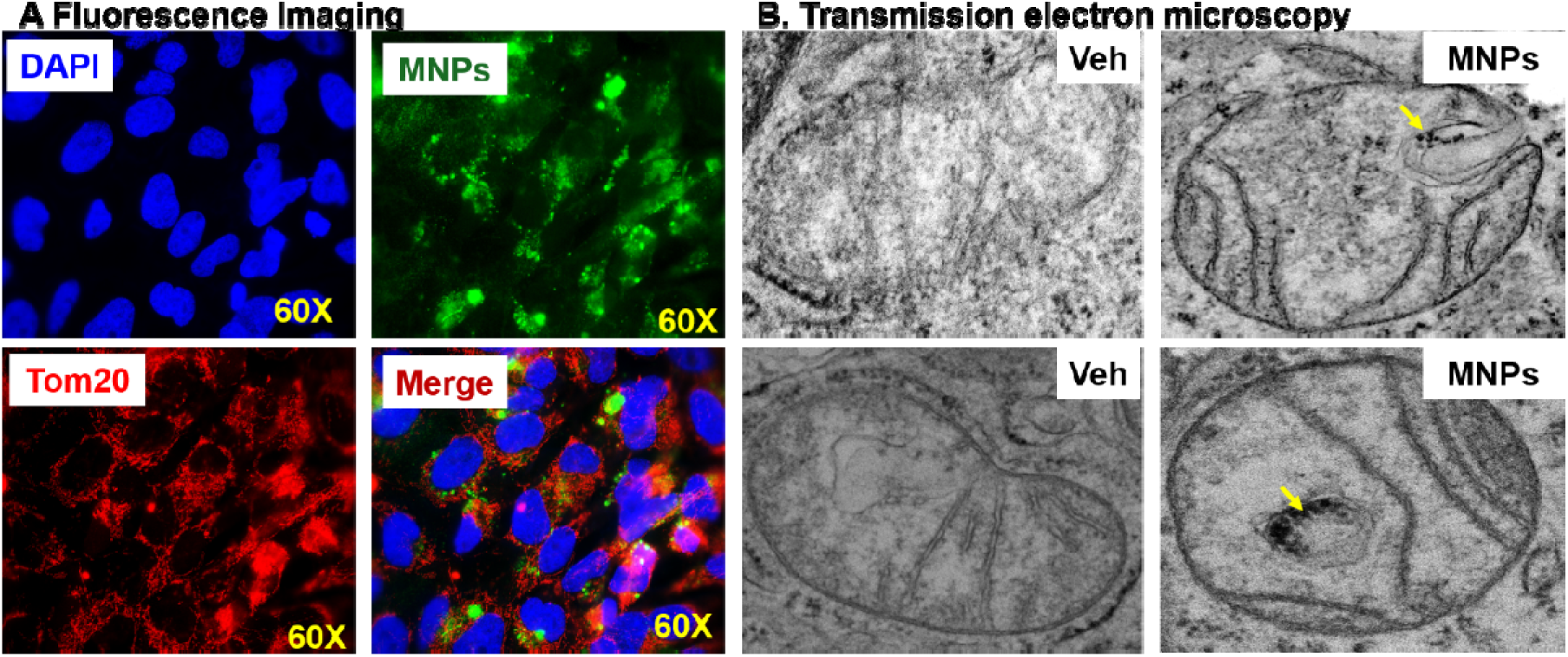
Representative images showing mitochondrial localization of MNPs in trophoblast cells. (A) Fluorescence microscopy images showing Green MNPs (25nm) co-localized with Tom20 Red (mitochondrial marker) and blue (nuclei). (B) TEM imaging shows MNP accumulation in mitochondria (yellow arrow) and structural changes (rounded) in the trophoblast after 96 hours of MNP exposure.

### MNP exposure disrupts cellular bioenergetics and mitochondrial respiration

To assess the impact of MNPs on mitochondrial function, we performed cellular bioenergetics analysis in trophoblast cells after 24 hours and 96 hours of MNPs exposure. We found that exposing trophoblasts to MNPs significantly impaired mitochondrial respiration and glycolytic function in a time-dependent manner. Vehicle treated cells maintained strong basal respiration, while MNP-treated cells exhibited a significant reduction in basal oxygen consumption rate (OCR) (p < 0.05) and decreased maximal respiration capacity post-FCCP treatment. ATP-linked respiration and spare respiratory capacity were also significantly reduced in MP-exposed cells, indicating impaired ATP production and diminished mitochondrial resilience to metabolic stress (Figure 5A, C). Measurements of extracellular acidification rate (ECAR, Figure 5 B) during mito stress tests revealed that MNP exposure caused a time-dependent decline in glycolytic activity, significantly lower than the control group (p < 0.05). Glycolytic capacity, measured after oligomycin injection, was also reduced, suggesting impaired glycolytic flux. The decline in both oxidative phosphorylation and glycolysis was more notable with longer MNPs exposure, linking it to mitochondrial dysfunction. Overall, these results indicate that MNPs exposure disrupts trophoblast energy metabolism.

**Figure 5.**
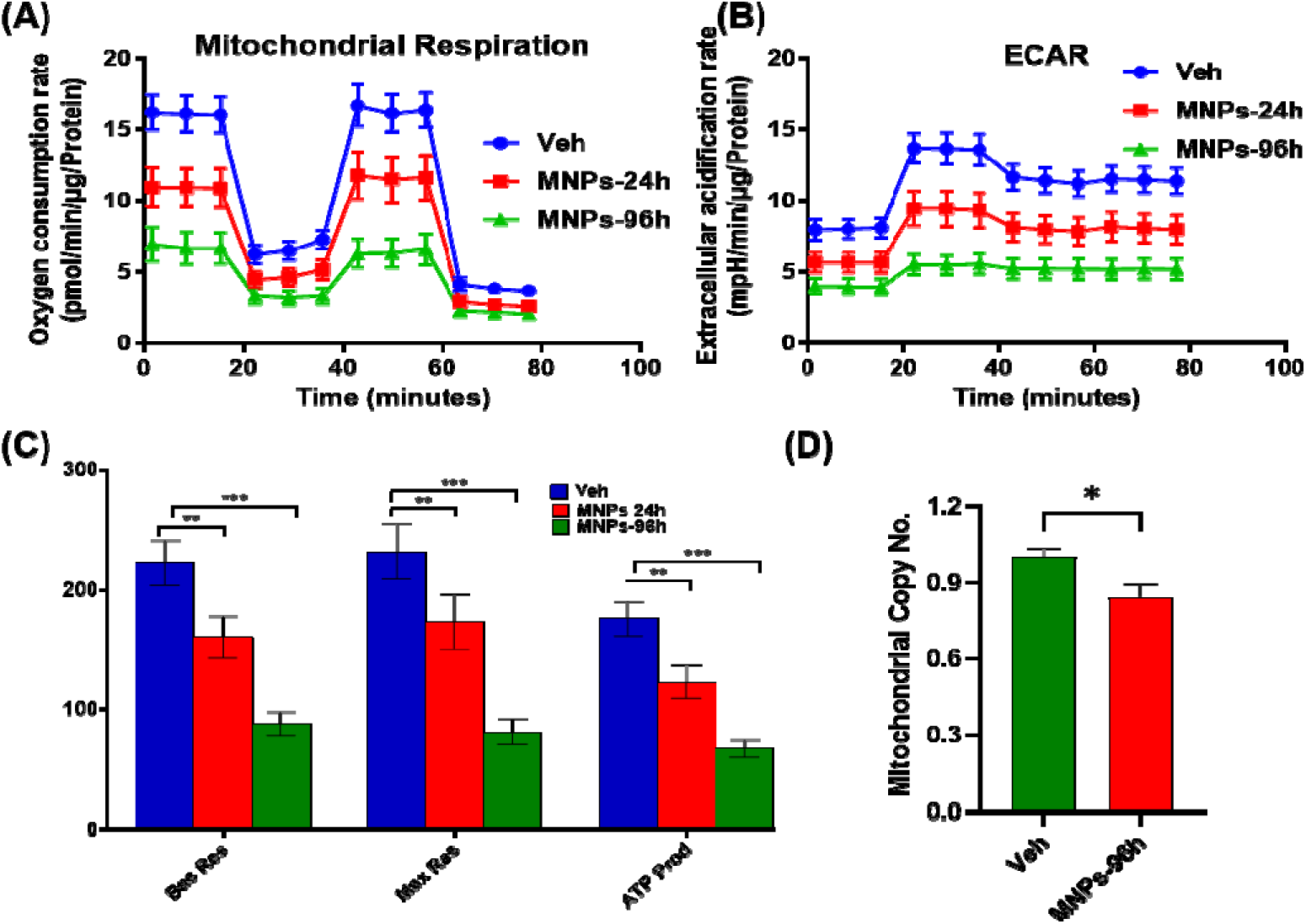
MNP uptake impair Mitochondrial Function in human Trophoblast cells (A) The oxygen consumption rate (OCR) profile, measured under basal conditions and following sequential treatment with oligomycin, FCCP, and rotenone/antimycin A, demonstrated impaired mitochondrial respiration in cells treated with MNPs compared to vehicle-treated human trophoblasts. (B) The extracellular acidification rate (ECAR) profile indicates altered glycolytic activity following MNP treatment. (C) Quantitative analysis of mitochondrial respiratory parameters, including basal respiration, maximal respiration, and ATP-linked respiration, showed a reduced mitochondrial functional capacity in the MNPs treated groups compared to vehicle controls. (D) Relative mitochondrial copy number in the MNPs and vehicle treatment group (data shown in fold change). Data are presented as mean ± SEM.; n = 6-12. Significance: *p < 0.05, **p < 0.01, ***p < 0.001 (one-way ANOVA for multiple groups, t-test for two groups).

### Mitochondrial copy number

Furthermore, we assessed mitochondrial copy number following MNP exposure. After 96 hours of treatment, a significant reduction in mitochondrial copy number was observed, suggesting that MNP intake disrupts mitochondrial homeostasis and impairs mitochondrial biogenesis and/or maintenance in trophoblast cells (Figure 5 D).

### MNPs Exposure Triggers Inflammatory Responses and Impairs Trophoblast Migration

To assess the effects of MNPs exposure on inflammatory signaling and trophoblast migration, we performed qPCR for mRNA expression of inflammasome markers (NLRP3, TLR9, IL-1β,) and in wound-healing assays in human trophoblast cells. MNP treatment significantly upregulated NLRP3 and TLR9, consistent with inflammasome activation, and markedly increased IL-1β expression, reflecting enhanced inflammatory signaling (Figure 6A). Vehicle-treated cells exhibited robust wound closure within 72 h, whereas MNPs-treated cells showed delayed closure with a persistent gap, indicating significantly impaired migratory capacity (Figure 6 B & C). Collectively, these findings demonstrate that exposure to MNPs activates inflammasome-mediated inflammatory pathways, thereby suppressing trophoblast motility, while supporting a potential role for MNPs in placental inflammation and growth.

**Figure 6.**
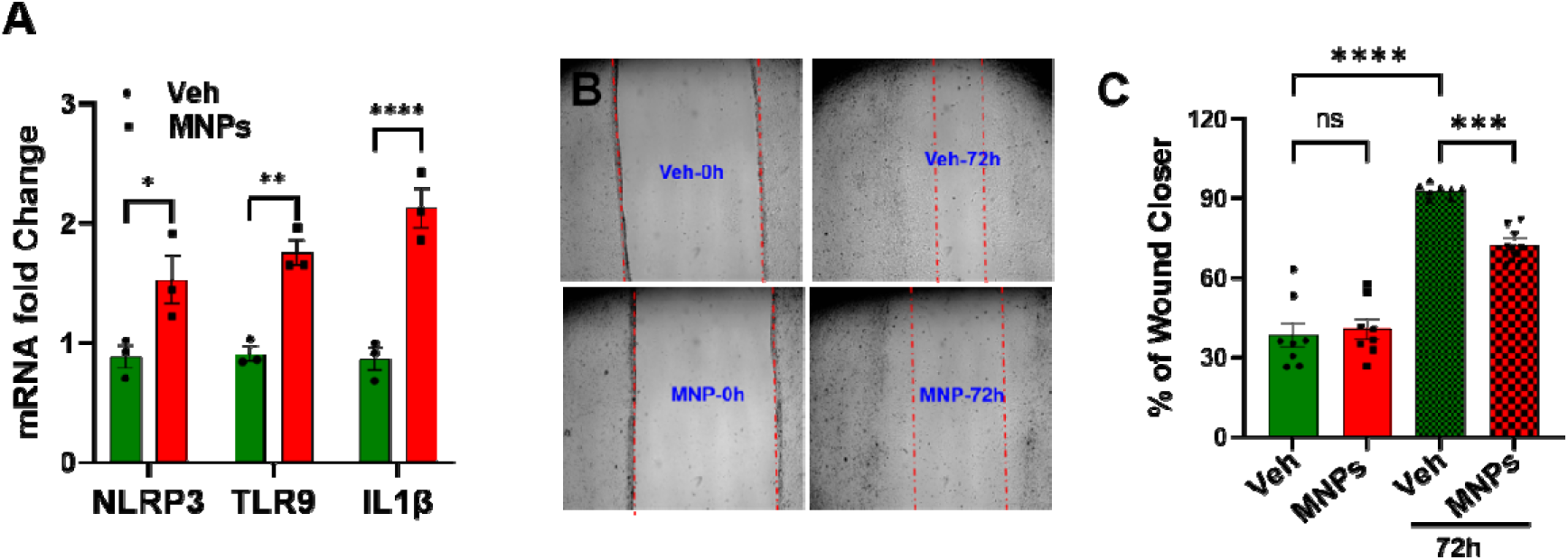
Human Trophoblast cells exposed to MNPS exhibit increased inflammation and reduced migratory capacity. (A) MNP exposure induces inflammatory responses (NLRP3, TLR9, and IL1β) in trophoblasts, as shown by qPCR analysis. (B) Wound closure assays demonstrated reduced healing in MNPs-treated trophoblasts compared to vehicle controls. N=3-4, *p < 0.05, **p < 0.01, ***p < 0.001, ****p < 0.0001 is considered significant. Data are presented as mean ± SEM.

## Discussion

Microplastics are ubiquitous environmental contaminants, and continuous human exposure occurs through air, water, and food, with increasing evidence linking micro and nano plastics (MNPs) to neurological [23,24], cardiovascular [25–27], and reproductive disorders in both males [28–30] and females [14,31,32]. Recent studies have identified diverse polymers, shapes, and sizes of MNPs in human tissues, including the placenta [4,5]. Both human and animal studies report placental accumulation of MNPs, which has been associated with adverse pregnancy outcomes such as preterm birth and miscarriage. In the present study, green-fluorescent MNPs were detected in multiple maternal organs (heart, lungs, kidneys, liver, and spleen) as well as in placental tissue, raising concerns about placental pathology and potential transplacental transfer with implications for fetal and neonatal health. Consistent with prior reports, our oral and intraperitoneal exposure models confirmed widespread tissue distribution of MNPs in pregnant mice, as visualized by IVIS imaging and organ-specific fluorescence analyses following exposure before breeding and throughout gestation.

This study aimed to elucidate the effects of MNP exposure on placental trophoblasts using an integrated in vivo and in vitro approach combined with fluorescence imaging and functional assays. Our findings demonstrate that MNPs are readily internalized by trophoblast cells, where they accumulate within intracellular and intra-organelle compartments, leading to pronounced alterations in cellular morphology and physiology. Notably, MNP accumulation induced significant mitochondrial structural and functional impairments alongside activation of inflammatory signaling pathways.

The placenta is a transient yet highly metabolically active organ that serves as the primary interface for maternal-fetal exchange, rendering it particularly vulnerable to environmental xenobiotics [33]. Its endocrine function and high energy demands rely heavily on tightly regulated mitochondrial activity, making placental tissue especially susceptible to mitochondrial stressors [34]. Fluorescence microscopy confirmed intracellular localization of polystyrene MNPs within trophoblasts, consistent with emerging evidence that MNPs can penetrate biological membranes and accumulate within intracellular compartments. Given their nanoscale size, MNPs may directly interact with mitochondria [35], thereby disrupting mitochondrial homeostasis [36]. Transmission electron microscopy revealed severe mitochondrial ultrastructural abnormalities, including intake of MNPs, swelling, disrupted cristae, vacuolization, and membrane damage, indicative of substantial mitochondrial injury. Importantly, the localization of MNPs within mitochondria suggests a direct contribution to mitochondrial oxidative stress. Supporting these observations, Seahorse bioenergetic analyses demonstrated significant impairments in oxygen consumption, including reduced basal respiration, ATP-linked respiration, and spare respiratory capacity, reflecting compromised oxidative phosphorylation and diminished metabolic efficiency in the trophoblast. The novel observation regarding mitochondrial uptake of MNPs opens exciting avenues for further research, necessitating validation across various cell types to deepen our understanding of this phenomenon.

Placental mitochondrial dysfunction has been closely associated with impaired trophoblast invasion, oxidative stress, abnormal spiral artery remodeling, and pregnancy complications such as preeclampsia. Excessive mitochondrial ROS production can further amplify inflammatory signaling and cellular damage, while reduced ATP availability may limit trophoblast adaptability to physiological stress, contributing to defective placentation [37–39]. In agreement with these mechanisms, our study also identified alterations in mitochondrial DNA copy number following MNP exposure, suggesting mitochondrial genomic instability and disrupted mitochondrial homeostasis.

In addition to mitochondrial impairment, quantitative real-time PCR analyses revealed dysregulation of inflammation-related genes in MNP-treated trophoblasts, indicating activation of cellular stress and inflammatory pathways. Mitochondrial dysfunction and ROS generation are tightly interconnected processes that can synergistically promote trophoblast injury [40]. These findings are particularly relevant to placental pathologies such as preeclampsia, where oxidative stress and mitochondrial abnormalities are central features [40,41].

Trophoblast proliferation, migration, and invasion are essential for normal placentation, spiral artery remodeling, and establishment of efficient maternal-fetal exchange [42–45]. While IL-1β plays a physiological role in regulating trophoblast invasion, we observed a significant increase in IL-1β expression following MNP exposure. Sustained elevation of IL-1β may promote chronic placental inflammation, dysregulated matrix remodeling, oxidative stress, and impaired trophoblast invasiveness. Consistent with this interpretation, wound-healing assays demonstrated significantly reduced migratory capacity in MNP-treated trophoblasts, a phenotype associated with defective placentation, fetal growth restriction, and preeclampsia.

A major strength of this study is the integration of multidisciplinary approaches, including in vivo exposure models, fluorescence imaging, ultrastructural analyses, bioenergetic profiling, and molecular assessments, providing a comprehensive evaluation of MNP-induced trophoblast dysfunction. However, limitations include the use of placental-derived trophoblast cell lines and a single exposure concentration, which may not fully reflect environmentally relevant human exposures. Future studies employing primary trophoblasts, varied exposure doses, and long-term in vivo models are warranted.

In conclusion, our findings demonstrate that MNP exposure induces significant trophoblast dysfunction characterized by intracellular accumulation, mitochondrial structural and bioenergetic impairment, and inflammatory activation. Collectively, these results identify mitochondrial dysfunction as a central mechanism underlying MNP-induced placental toxicity and highlight MNPs as emerging environmental risk factors for placental disorders and adverse pregnancy outcomes.

## Contribution

PKD conceptualized, designed, and supervised the study. PV, SD, and PKD performed the experiments, analyzed the data, and drafted the manuscript. VR provided technical assistance with the experimental procedures. SD, YW, PC, and DB contributed critical intellectual insights and provided valuable feedback during the study. All authors reviewed, edited, and approved the final version of the manuscript.

## Declaration of generative AI and technologies used

The author(s) used Grammarly to enhance clarity and correct syntax error.

## Acknowledgments

We would like to thank the UAB Small Animal Imaging Facility (SAIF), the High-Resolution Imaging Facility (HRIF), Dr. Pulin Che’s lab for using fluorescence microscopy, and Dr. Yajing Wang’s lab for the use of Agilent Seahorse XF system in this study at the University of Alabama at Birmingham, AL.

## Data availability

Data will be made available on reasonable request.

## References

[1] G. Zuri, A. Karanasiou, S. Lacorte, Microplastics: Human exposure assessment through air, water, and food, Environment International 179 (2023) 108150. 10.1016/j.envint.2023.108150.

[2] F. Jahedi, N. Jaafarzadeh Haghighi Fard, Micro- and nanoplastic toxicity in humans: Exposure pathways, cellular effects, and mitigation strategies, Toxicol Rep 14 (2025) 102043. 10.1016/j.toxrep.2025.102043.

[3] Q. Yang, Y. Peng, X. Wu, X. Cao, P. Zhang, Z. Liang, J. Zhang, Y. Zhang, P. Gao, Y. Fu, P. Liu, Z. Cao, T. Ding, Microplastics in human skeletal tissues: Presence, distribution and health implications, Environ Int 196 (2025) 109316. 10.1016/j.envint.2025.109316.

[4] M.A. Garcia, R. Liu, A. Nihart, E. El Hayek, E. Castillo, E.R. Barrozo, M.A. Suter, B. Bleske, J. Scott, K. Forsythe, J. Gonzalez-Estrella, K.M. Aagaard, M.J. Campen, Quantitation and identification of microplastics accumulation in human placental specimens using pyrolysis gas chromatography mass spectrometry, Toxicol Sci 199 (2024) 81–88. 10.1093/toxsci/kfae021.

[5] A. Ragusa, A. Svelato, C. Santacroce, P. Catalano, V. Notarstefano, O. Carnevali, F. Papa, M.C.A. Rongioletti, F. Baiocco, S. Draghi, E. D’Amore, D. Rinaldo, M. Matta, E. Giorgini, Plasticenta: First evidence of microplastics in human placenta, Environ Int 146 (2021) 106274. 10.1016/j.envint.2020.106274.

[6] O. Khaybullina, P. Sarkhel, O.M. Keinänen, Tracking of [14C]Polystyrene Nanoplastics in Pregnant Mice, Adv Sci (Weinh) 13 (2026) e23995. 10.1002/advs.202523995.

[7] C. Wang, H. Chang, H. Wang, H. Li, S. Ding, F. Ren, Exposure to microplastics during pregnancy and fetal liver function, Ecotoxicology and Environmental Safety 294 (2025) 118099. 10.1016/j.ecoenv.2025.118099.

[8] M. Jochum, M. Garcia, A. Hammerquist, J. Howell, M. Stanford, R. Liu, M. Olewine, E.E. Hayek, E. Phan, L. Showalter, C. Shope, M. Suter, M. Campen, K. Aagaard, E. Barrozo, Elevated Micro- and Nanoplastics Detected in Preterm Human Placentae, Res Sq (2025) rs.3.rs-5903715. 10.21203/rs.3.rs-5903715/v1.

[9] R. Zhang, Y.I. Zhou, X. Zhao, Q. Chen, X. Li, Y. Chen, J. Xi, R. Yang, B.O. Xie, Y. Yang, T. Zhai, Y. Meng, L. Chen, Z. Yan, Y.A. Qi, F. Xiang, W. Zheng, S. Jiang, T. Cao, Y. Wang, Y. Zhou, S. Yan, Identification and analysis of microplastics in main organs of human fetus, Ecotoxicology and Environmental Safety 314 (2026) 120078. 10.1016/j.ecoenv.2026.120078.

[10] C.Q.Y. Yong, S. Valiyaveettil, B.L. Tang, Toxicity of Microplastics and Nanoplastics in Mammalian Systems, Int J Environ Res Public Health 17 (2020) 1509. 10.3390/ijerph17051509.

[11] B. Li, Y. Ding, X. Cheng, D. Sheng, Z. Xu, Q. Rong, Y. Wu, H. Zhao, X. Ji, Y. Zhang, Polyethylene microplastics affect the distribution of gut microbiota and inflammation development in mice, Chemosphere 244 (2020) 125492. 10.1016/j.chemosphere.2019.125492.

[12] S. Wan, X. Wang, W. Chen, M. Wang, J. Zhao, Z. Xu, R. Wang, C. Mi, Z. Zheng, H. Zhang, Exposure to high dose of polystyrene nanoplastics causes trophoblast cell apoptosis and induces miscarriage, Part Fibre Toxicol 21 (2024) 13. 10.1186/s12989-024-00574-w.

[13] P. Wang, G. Li, H. Shou, X. Huang, H. Xu, S. Zhang, H. Zhu, Microplastic abundance in placental chorionic villi detected by pyrolysis-gas chromatography/mass spectrometry in cases of spontaneous miscarriage during early pregnancy, eBioMedicine 120 (2025) 105918. 10.1016/j.ebiom.2025.105918.

[14] Z. Liu, Q. Zhuan, L. Zhang, L. Meng, X. Fu, Y. Hou, Polystyrene microplastics induced female reproductive toxicity in mice, Journal of Hazardous Materials 424 (2022) 127629. 10.1016/j.jhazmat.2021.127629.

[15] Y. Wang, S. Zhao, Cell Types of the Placenta, in: Vascular Biology of the Placenta, Morgan & Claypool Life Sciences, 2010. https://www.ncbi.nlm.nih.gov/books/NBK53245/ (accessed June 21, 2026).

[16] M. Knöfler, S. Haider, L. Saleh, J. Pollheimer, T.K.J.B. Gamage, J. James, Human placenta and trophoblast development: key molecular mechanisms and model systems, Cell Mol Life Sci 76 (2019) 3479–3496. 10.1007/s00018-019-03104-6.

[17] S. Jinesh, P. Aditi, Health Implications of Microplastic Exposure in Pregnancy and Early Childhood: A Systematic Review, Int J Womens Health 17 (2025) 2805–2818. 10.2147/IJWH.S497366.

[18] M.C. Bucknor, B.A. Keating, V.X. Han, B.S. Gloss, P. Dey, N. Aryamanesh, L.L. Marshall, M.E. Graham, R. Dissanayake, X. Lau, S. Patel, S.P. Petkova, A. Gururajan, R.C. Dale, M.J. Hofer, Cumulative pregnancy and postnatal environmental exposures impact social behaviour in male mice associated with epigenetic, ribosomal, and immune dysregulation, (2025) 2025.03.19.644073. 10.1101/2025.03.19.644073.

[19] R. Naha, A. Anees, S. Chakrabarty, P.S. Naik, M. Pandove, D. Pandey, K. Satyamoorthy, Placental mitochondrial DNA mutations and copy numbers in intrauterine growth restricted (IUGR) pregnancy, Mitochondrion 55 (2020) 85–94. 10.1016/j.mito.2020.08.008.

[20] R.D. Williamson, F.P. McCarthy, A.S. Khashan, A. Totorika, L.C. Kenny, C. McCarthy, Exploring the role of mitochondrial dysfunction in the pathophysiology of pre-eclampsia, Pregnancy Hypertens 13 (2018) 248–253. 10.1016/j.preghy.2018.06.012.

[21] V. Rajasekaran, S. Dubey, T. Kaushik, A.G. Neuenschwander, N.-S. Rajasekaran, D.E. Berkowitz, P.K. Dubey, Mitochondrial Heterogeneity in a Patient with Preeclampsia, Free Radical Biology and Medicine (2026). 10.1016/j.freeradbiomed.2026.02.025.

[22] Rajasekaran, V., et al., Mitochondrial heterogeneity in a patient with preeclampsia. Free Radical Biology and Medicine, 2026. 247: p. 315–318. - Google Search, (n.d.). https://www.google.com/search?q=Rajasekaran%2C+V.%2C+et+al.%2C+Mitochondrial+heterogeneity+in+a+patient+with+preeclampsia.+Free+Radical+Biology+and+Medicine%2C+2026.+247%3A+p.+315-318. (accessed June 11, 2026).

[23] A.J. Nihart, M.A. Garcia, E. El Hayek, R. Liu, M. Olewine, J.D. Kingston, E.F. Castillo, R.R. Gullapalli, T. Howard, B. Bleske, J. Scott, J. Gonzalez-Estrella, J.M. Gross, M. Spilde, N.L. Adolphi, D.F. Gallego, H.S. Jarrell, G. Dvorscak, M.E. Zuluaga-Ruiz, A.B. West, M.J. Campen, Bioaccumulation of microplastics in decedent human brains, Nat Med 31 (2025) 1114–1119. 10.1038/s41591-024-03453-1.

[24] Nanoplastics in the human brain and their change in abundance over time, Nat Med 31 (2025) 1077–1078. 10.1038/s41591-025-03571-4.

[25] R. Marfella, F. Prattichizzo, C. Sardu, G. Fulgenzi, L. Graciotti, T. Spadoni, N. D’Onofrio, L. Scisciola, R.L. Grotta, C. Frigé, V. Pellegrini, M. Municinò, M. Siniscalchi, F. Spinetti, G. Vigliotti, C. Vecchione, A. Carrizzo, G. Accarino, A. Squillante, G. Spaziano, D. Mirra, R. Esposito, S. Altieri, G. Falco, A. Fenti, S. Galoppo, S. Canzano, F.C. Sasso, G. Matacchione, F. Olivieri, F. Ferraraccio, I. Panarese, P. Paolisso, E. Barbato, C. Lubritto, M.L. Balestrieri, C. Mauro, A.E. Caballero, S. Rajagopalan, A. Ceriello, B. D’Agostino, P. Iovino, G. Paolisso, Microplastics and Nanoplastics in Atheromas and Cardiovascular Events, New England Journal of Medicine 390 (2024) 900–910. 10.1056/NEJMoa2309822.

[26] Y.-H. Lee, C.-M. Zheng, Y.-J. Wang, Y.-L. Wang, H.-W. Chiu, Effects of microplastics and nanoplastics on the kidney and cardiovascular system, Nat Rev Nephrol 21 (2025) 585–596. 10.1038/s41581-025-00971-0.

[27] A. Goldsworthy, L.A. O’Callaghan, C. Blum, J. Horobin, L. Tajouri, M. Olsen, N. Van Der Bruggen, S. McKirdy, R. Alghafri, O. Tronstad, J. Suen, J.F. Fraser, Micro-nanoplastic induced cardiovascular disease and dysfunction: a scoping review, J Expo Sci Environ Epidemiol 35 (2025) 746–769. 10.1038/s41370-025-00766-2.

[28] Y.-Y. Hwang, Q.-T. Tsen, J. Felim, R.-F. Shiu, F. Kong, Z.-L. Kong, D.-F. Hwang, Probiotics as a therapeutic approach to alleviate reproductive harm from polystyrene microplastics in male rats, Sci Rep 15 (2025) 34783. 10.1038/s41598-025-18550-5.

[29] J. Codrington, A.A. Varnum, L. Hildebrandt, D. Pröfrock, J. Bidhan, K. Khodamoradi, A.-L. Höhme, M. Held, A. Evans, D. Velasquez, C.C. Yarborough, B. Ghane-Motlagh, A. Agarwal, J. Achua, E. Pozzi, F. Mesquita, F. Petrella, D. Miller, R. Ramasamy, Detection of microplastics in the human penis, Int J Impot Res 37 (2025) 377–383. 10.1038/s41443-024-00930-6.

[30] A.H.A. Alsenousy, A.H.Y. Khalaf, H.Z. Ibrahim, M.A. Kamel, M.I. Yousef, Impact of polystyrene microplastic exposure at low doses on male fertility: an experimental study in rats, Sci Rep 16 (2026) 7474. 10.1038/s41598-026-38385-y.

[31] A. Haddadi, K. Kessabi, S. Boughammoura, M.B. Rhouma, R. Mlouka, M. Banni, I. Messaoudi, Exposure to microplastics leads to a defective ovarian function and change in cytoskeleton protein expression in rat, Environ Sci Pollut Res 29 (2022) 34594–34606. 10.1007/s11356-021-18218-3.

[32] J. Hou, Z. Lei, L. Cui, Y. Hou, L. Yang, R. An, Q. Wang, S. Li, H. Zhang, L. Zhang, Polystyrene microplastics lead to pyroptosis and apoptosis of ovarian granulosa cells via NLRP3/Caspase-1 signaling pathway in rats, Ecotoxicology and Environmental Safety 212 (2021) 112012. 10.1016/j.ecoenv.2021.112012.

[33] A.F. Sobral, A. Cunha, I. Costa, M. Silva-Carvalho, R. Silva, D.J. Barbosa, Environmental Xenobiotics and Epigenetic Modifications: Implications for Human Health and Disease, Journal of Xenobiotics 15 (2025) 118. 10.3390/jox15040118.

[34] F. Jahan, G. Vasam, A.E. Green, S.A. Bainbridge, K.J. Menzies, Placental Mitochondrial Function and Dysfunction in Preeclampsia, Int J Mol Sci 24 (2023) 4177. 10.3390/ijms24044177.

[35] T.-H. Chen, I.-T. Chu, R.-Y. Chang, H.-C. Wang, C.-J. Chung, T.-H. Tsai, C.-K. Chou, Microplastics induce mitochondrial dysfunction and accelerate cardiovascular pathogenesis, Arch Toxicol 100 (2026) 1699–1711. 10.1007/s00204-026-04327-w.

[36] A.F. Hernández, M. Lacasaña, A.M. Tsatsakis, A.O. Docea, Cellular and Molecular Mechanisms of Micro- and Nanoplastics Driving Adverse Human Health Effects, Toxics 13 (2025) 921. 10.3390/toxics13110921.

[37] I. Vornic, V. Buciu, C.G. Furau, P.N. Gaje, R.A. Ceausu, C.-S. Dumitru, A.C. Barb, D. Novacescu, A.A. Cumpanas, S.C. Latcu, T.G. Cut, F. Zara, Oxidative Stress and Placental Pathogenesis: A Contemporary Overview of Potential Biomarkers and Emerging Therapeutics, Int J Mol Sci 25 (2024) 12195. 10.3390/ijms252212195.

[38] J.J. Fisher, L.A. Bartho, A.V. Perkins, O.J. Holland, Placental mitochondria and reactive oxygen species in the physiology and pathophysiology of pregnancy, Clin Exp Pharmacol Physiol 47 (2020) 176–184. 10.1111/1440-1681.13172.

[39] J. Long, Y. Huang, G. Wang, Z. Tang, Y. Shan, S. Shen, X. Ni, Mitochondrial ROS Accumulation Contributes to Maternal Hypertension and Impaired Remodeling of Spiral Artery but Not IUGR in a Rat PE Model Caused by Maternal Glucocorticoid Exposure, Antioxidants (Basel) 12 (2023) 987. 10.3390/antiox12050987.

[40] I. Vasilaki, A. Potiris, E. Moustakli, D. Mavrogianni, N. Daponte, T. Karampitsakos, A. Kozonis, K. Louis, C. Messini, T. Grigoriadis, E. Domali, S. Stavros, Linking Oxidative Stress to Placental Dysfunction: The Key Role of Mitochondria in Trophoblast Function, Med Sci (Basel) 14 (2026) 53. 10.3390/medsci14010053.

[41] F. Khadir, Z. Rahimi, A. Ghanbarpour, A. Vaisi-Raygani, Nrf2 rs6721961 and Oxidative Stress in Preeclampsia: Association with the Risk of Preeclampsia and Early-Onset Preeclampsia, Int J Mol Cell Med 11 (2022) 127–136. 10.22088/IJMCM.BUMS.11.2.127.

[42] S.J. Renaud, K. Kubota, M.A.K. Rumi, M.J. Soares, The FOS transcription factor family differentially controls trophoblast migration and invasion, J Biol Chem 289 (2014) 5025–5039. 10.1074/jbc.M113.523746.

[43] P. Velicky, M. Knöfler, J. Pollheimer, Function and control of human invasive trophoblast subtypes: Intrinsic vs. maternal control, Cell Adh Migr 10 (2015) 154–162. 10.1080/19336918.2015.1089376.

[44] C.-C. Huang, Y.-W. Hsueh, C.-W. Chang, H.-C. Hsu, T.-C. Yang, W.-C. Lin, H.-M. Chang, Establishment of the fetal-maternal interface: developmental events in human implantation and placentation, Front. Cell Dev. Biol. 11 (2023). 10.3389/fcell.2023.1200330.

[45] X. Gan, H. Liu, D. Chen, Z. Liu, Q. Lu, X. Lai, H. Hou, M. Zhang, J.Y. Zhang, Y. Duan, S. Lu, M. Chen, G.E. Lash, F. Ning, Interleukin-1 beta signals through the ERK signalling pathway to modulate human placental trophoblast migration and invasion in the first trimester of pregnancy, Placenta 151 (2024) 67–78. 10.1016/j.placenta.2024.04.010.

